# Helical ensembles out-perform ideal helices in Molecular Replacement

**DOI:** 10.1101/2020.06.16.154690

**Authors:** Filomeno Sánchez Rodríguez, Adam J. Simpkin, Owen R. Davies, Ronan M. Keegan, Daniel J. Rigden

**Affiliations:** Institute of Structural, Molecular and Integrative Biology, University of Liverpool, Liverpool L69 7ZB, England; Life Science, Diamond Light Source, Harwell Science and Innovation Campus, Didcot, Oxfordshire OX11 0DE, England; Institute for Cell and Molecular Biosciences, Newcastle University, Framlington Place, Newcastle upon Tyne NE2 4HH, England; UKRI-STFC, Rutherford Appleton Laboratory, Research Complex at Harwell, Didcot OX11 0FA, England

## Abstract

The conventional approach in molecular replacement (MR) is the use of a related structure as a search model. However, this is not always possible as the availability of such structures can be scarce for poorly characterised families of proteins. In these cases, alternative approaches can be explored, such as the use of small ideal fragments that share high albeit local structural similarity with the unknown protein. Earlier versions of *AMPLE* enabled the trialling of a library of ideal helices, which worked well for largely helical proteins at suitable resolution. Here we explore the performance of libraries of helical ensembles created by clustering helical segments. The impacts of different B-factor treatments and different degrees of structural heterogeneity are explored. We observed a 30% increase in the number of solutions obtained by *AMPLE* when using this new set of ensembles compared to performance with ideal helices. The boost of performance was notable across three different folds: transmembrane, globular and coiled-coil structures. Furthermore, the increased effectiveness of these ensembles was coupled to a reduction of the time required by *AMPLE* to reach a solution. *AMPLE* users can now take full advantage of this new library of search models by activating the “helical ensembles” mode.

## 1. Introduction

X-ray crystallography is the most prevalent technique for protein structure determination (Berman et al., 2002) but the phase problem remains one of its most challenging aspects. Molecular replacement (MR) is often the method of choice to obtain the missing phase information, primarily because of its speed and high potential for automation (Evans & McCoy, 2007). This approach relies on replacing the missing experimental phases of the unknown structure with the calculated phases of a similar solved structure (Rossmann, 1990), positioned appropriately in the unit cell. Thus, the more structurally similar the search model is to the unknown structure, the more probable MR will have a positive outcome. In non-trivial MR cases, the availability and detection of suitable search models can be a key limitation and alternative routes need to be explored. One such route is the use of small fragments such as α-helices. This was first proposed as an efficient phasing technique to solve cases where high resolution scattering data was available (Yao et al., 2002). Since most proteins contain α-helices or *β* -strands as secondary structure elements, such standardised fragments are often a valid approximation to elements of the unknown structure. Nevertheless, the use of these search models has the intrinsic difficulty of low signal to noise ratio as they only represent a small fraction of the overall structure. Such complications were addressed by the development of *ARCIMBOLDO* (Rodríguez et al., 2009), which combined the use of *PHASER* (McCoy et al., 2007) for the accurate placement of several small ideal fragments with *SHELXE* (Thorn & Sheldrick, 2013) for electron density map modification and chain autotracing. The use of this sophisticated approach enabled the solution of increasingly challenging structures with lower resolutions -between 2.5 Å and 3 Å- and larger numbers of residues in the asymmetric unit -between 400 and 500 residues-. Additionally, this approach proved successful on the solution of a variety of folds, such as coiled-coils (Caballero et al., 2018), which are notoriously difficult to solve through MR as they often suffer from different crystallographic data pathologies such as anisotropic diffraction and translational non-crystallographic symmetry. Further developments in fragment-based MR came with *FRAGON* (Jenkins, 2018). This tool achieves high success rates among high resolution structures while having a relatively low consumption of computational resources by taking advantage of *PHASER’S* ability to place small search fragments and *ACORN’s* (J. X. Yao et al., 2006) sophisticated scoring algorithm for density modification.

*AMPLE* first made use of such standard fragments as a baseline to assess the performance of *ab initio* modelling decoys as search models for the solution of the aforementioned challenging coiled-coil structures (Thomas et al., 2015). This small library of eight ideal helices, with lengths ranging from 5 to 40 residues, proved to be surprisingly effective as more than half of the structures under study could be solved, thus revealing a fast and simple MR approach. Later studies suggested the efficacy of *AMPLE’s* ideal helices extends to different fold types, such as α-helical transmembrane proteins (Thomas et al., 2017). However, despite the relative success observed when using these search models, limitations in the use of these helices as search models could be observed, as revealed by the scarce presence of solved structures with more than 300 residues in the asymmetric unit and scattering data with resolutions of 2 Å or worse. Nevertheless, such limitations are to be expected, as idealised fragments cannot capture in full the details of biologically active protein folds, which are a result of a combination of biochemical interactions that will cause a series of distortions in the local fold of the structure (Koga et al., 2012). However, several tools have been developed in the field of fragment-based MR aiming on this and other limitations intrinsic to fragment search models, such as *BORGES* (Sammito et al., 2013), which makes use of multiple fragments to expand the size of the search model into recurring tertiary structure motifs, *ALEPH* (Medina et al., 2020), which mines structures available at databases to create libraries of fragments that can be used as search models, or *ALIXE* (Millán et al., 2020), which combines phase information from partial solutions originating from these correctly placed fragment search models.

Here we explore new ways to increase the efficacy of helical fragments as search models in *AMPLE,* by making use of ensemble search models. Ensembles of multiple search models created through structural alignment have a long history of out-performing their individual component structures (Bibby et al., 2012; Chen et al., 2000; Keegan et al., 2018; Leahy et al., 1992; Pieper et al., 1998; Rigden et al., 2002; Simpkin et al., 2020). With ensemble search models, the structural variability of the members in the ensemble can be used to statistically weight sets of structure factors (Read, 2001). We observe that a new set of 64 helical ensembles created by mining the currently available structures at the PDB (Berman et al., 2002) and superimposing detected helices, significantly outperforms the original set of single model ideal helices as search models. Additionally, we compare MR success rates for ensembles with different levels of structural heterogeneity and subject to different B-factor treatments. Compared with the original set of ideal helices, we registered a 30% increase in the number of solutions when using the new library of helical ensembles, an improvement which is consistent across transmembrane, globular and coiled-coil folds. This increase in the number of MR successes is coupled with a decrease in the time elapsed before reaching the first solution for a given structure when using a minimal subset of 12 ensembles.

## 2. Methods

### 2.1 Dataset selection

The test set of globular and transmembrane structures was selected by first retrieving a list of all the X-ray PDB entries with a resolution between 2.0 and 3.0 Å from which chain lengths between 100 and 700 residues were selected. Transmembrane protein entries meeting these requirements were determined by reference to the PDBTM register (Kozma et al., 2013). Those structures within the globular set annotated as coiled-coils by the SCOP classification (Andreeva et al., 2020) were split into a coiled-coil set. For the resulting three datasets, the sequences of all structures were clustered using *CD-HIT* (Fu et al., 2012), which also identified a representative sequence for each cluster using an identity cutoff of 20%. In order to remove redundancy among the structures of the three sets, only the structures of the representative sequences were kept in the dataset. *PHASER’S* expected log-likelihood gain (eLLG) (Oeffner et al., 2018) reflects the log-likelihood gain on intensity (LLGI) expected from a correctly placed model, and it can be used for the purpose of assessing the difficulty of solving a structure through MR with a given search model. In order to form a set of structures representative of different ranges of difficulty possible to encounter while solving different MR cases, the structures were organised into four bins according to their eLLG values obtained with a 40 residue-long poly-alanine ideal helix as a search model and an expected RMS value of 0.1. The ranges of these bins were set using the values indicated in *PHASER’S* guidelines (Oeffner et al., 2018): above 64, between 64-49, between 49-36, and between 36-25. This resulted in the final selection of 34 transmembrane (Supplementary Table 1), 31 globular (Supplementary Table 2) structures and 18 coiled-coils. Due to the reduced number of coiled-coil structures found with this procedure, we extended this dataset to include 28 structures used in previous studies (Thomas et al., 2020), and allowed for increasingly difficult structures having eLLG values lower than 25. This resulted in the final set of 46 coiled-coils used in this study (Supplementary Table 3), with structures outside the 2 - 3 Å resolution range. This includes structures 3HFE, 4DZK, 1G1J, 1Y66, 3Q8T and 2EFR with resolutions better than 2 Å, the highest resolution being 3HFE at 1.7 Å, and structures 4U5T, 6GBR, 3MQB, 6BRI and 4GKW with resolutions worse than 3 Å, the lowest resolution being 4U5T and 4GKW at 3.30 Å.

### 2.2 Creation of the helical ensembles

In order to assess if the structural divergence between the models that form an ensemble has effects over its effectiveness as a search model, both low divergence (homogeneous) and high divergence (heterogeneous) ensembles were created. This was done by first performing a secondary structure search of all structures in the PDB (Berman et al., 2002) with *GESAMT* (Krissinel, 2012), using each of the eight ideal helices first used in *AMPLE* (Thomas et al., 2015) as search queries. The top four hits within an RMSD of 0.5 Å across all C_α_ atoms of the helix in the case of homogeneous ensembles, and between 0.5 - 1.0 Å for the heterogeneous ensembles were then structurally aligned with the original ideal helix to generate ensembles in the library (Supplementary Table 4). Finally, all side chains present in the models were removed in order to create polyalanine helices.

### 2.3 B-Factor treatments of the helical ensembles

B-factors play an important role in *PHASER’S* algorithm, as it bases the calculation of the structure factors on the normalised values of the B-factors of the atoms present in the search model. In order to assess whether B-factor adjustments reflecting the variability across models in different regions of the ensemble could increase the effectiveness of the search model during MR searches, the residue B-factors of the helical ensembles were modified using four different strategies (Figure 1). First, in the B-factor treatment 1, the native B-factors observed in the crystal structure were kept unmodified. In the case of B-factor treatment 2, B-factors were modified along a gradient, with values starting at a value of 10.0 Å^2^ on the two residues located at the centre of the helix -central residue numbers were rounded up in the case of even helical sizes- and increasing up to a limit of 90.0 Å^2^ towards the extremes. Uniform steps were used to increase B-factors between residues, and all the atoms of each residue were assigned the same value. In treatments number 3 and 4, B-factors were modified according to the structural variance observed across the models of the ensemble at each residue position. To do this, distances between all the C_α_ of the residues at equivalent positions across the models of the ensemble were measured. In the case of treatment number 3, the mean distance between each member of the ensemble in turn and the rest of its equivalent residues was measured, and B-factors were set for all atoms at each residue with values of 80.0 Å^2^ if the distance was higher than 0.8 Å, a value of 10.0 Å^2^ if the average distance was lower than 0.1 Å and a B-factor resulting from the following equation:

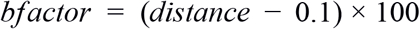

was used for any other value (see Figure 2 for detailed view). Finally, in the case of treatment number 4, the same B-factor was set at each position across the residues of all five models of the ensemble. For each position of the ensemble, the average distance between all equivalent residues was measured and the B-factor set as stated on treatment 3.

**Figure 1.**
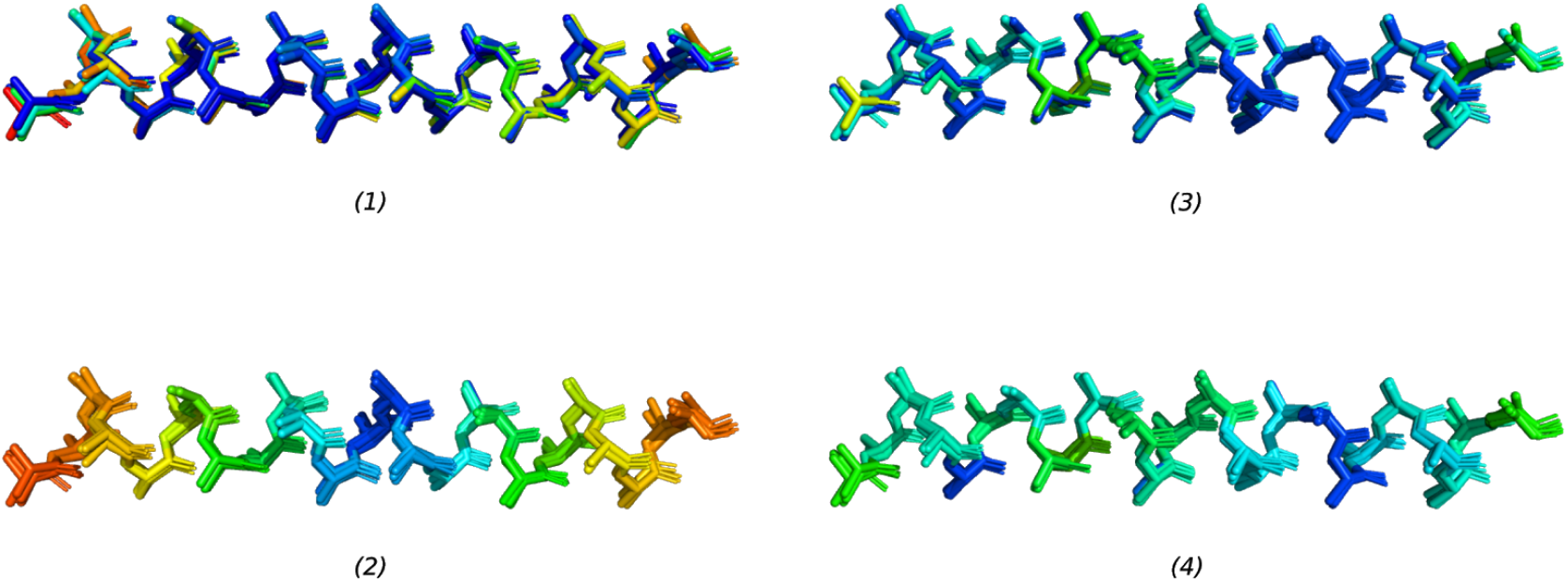
Depiction of the strategies used for the modification of B-factors of the ensembles. Residue colours correlate with their assigned B-factor, and follows a scale from blue -lower- to red -higher- going through green. All the treatments are illustrated using the 25 residue long homogeneous ensemble. Treatment 1: Native B-factors kept unmodified as in the native crystal structure. Treatment 2: Gradient of B-factors is created through the helical ensemble. Treatment 3: Each residue has a B-factor proportional to its mean distance with the rest of equivalent residues at the other four models. Treatment 4: Each residue on the ensemble is set with a B-factor proportional to the average distance of all the models in that position.

**Figure 2.**
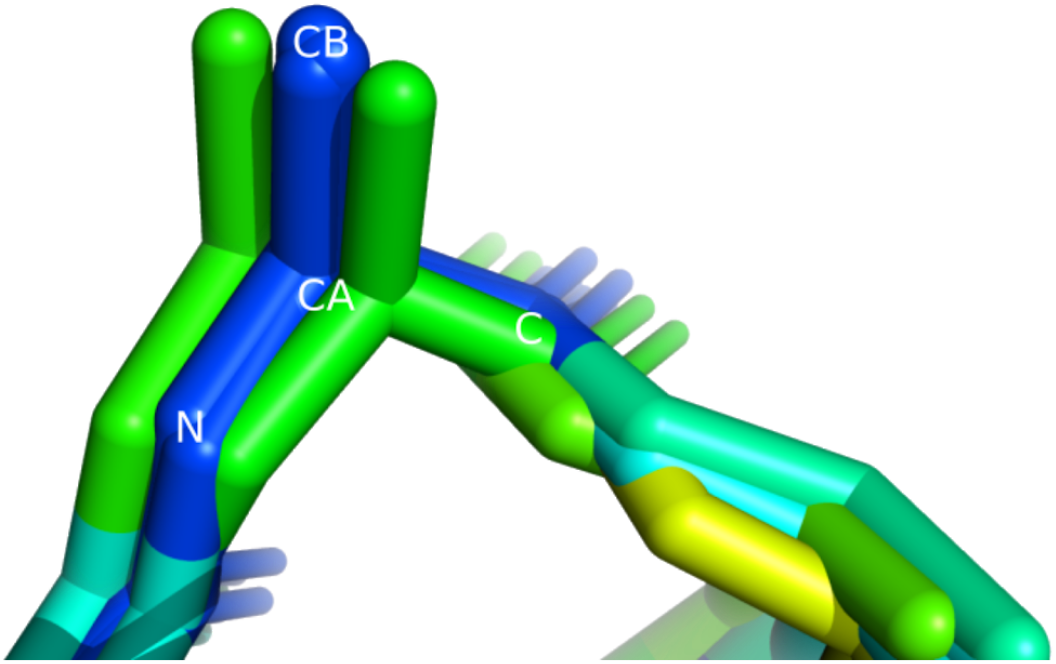
Detailed view of the 8th alanine residue of the 25 residue long homogenous ensemble where the B-factor has been modified as in treatment 3: the B-factor value is proportional to the average distance between each residue and the other residues at the equivalent positions of the other four models. Thus it is higher - green rather than blue - for outlying residues at this position in the ensemble.

### 2.4 Molecular replacement

Each structure of the dataset was trialled using two independent *AMPLE* MR runs, one using the original set of ideal helices - ideal helix mode - and another one using the new helical ensembles as an alternative -helical ensemble mode-. *AMPLE* makes use of the pipeline *MrBUMP* (Keegan et al., 2018) which in turn uses *PHASER* (McCoy et al., 2007) for molecular replacement, *REFMAC5* (Vagin et al., 2004) for refinement, and *SHELXE* (Thorn & Sheldrick, 2013) for density modification and c-apha tracing. For the purpose of this study, the *PHASER* kill-time value was set to 24 hours and in the case of coiled-coil folds, translational non-crystallographic symmetry corrections in *PHASER* were turned off. *PHASER* calculated a variance-r.m.s. (VRMS) parameter for each of the input ensembles to optimize the calculation of its log-likelihood gain score (LLG) and improve the chances of picking out a correct solution (Oeffner et al., 2013). This gave calculated VRMS values between 0.2 and 1.5 depending on the ensemble.

To determine the success of the molecular replacement attempts, at first it was determined if the search model had been placed correctly by calculating the correlation coefficient between the electron density map of the deposited experimental data and the map of the placed model (MapCC), using *phenix.get_cc_mtz_mtz* from the *PHENIX* suite. Search models were then considered correctly placed if this coefficient reached at least 0.2. Nevertheless, a correct placement does not always imply that it will be possible to trace the model and solve the unknown structure. In this particular aspect, the *SHELXE* correlation coefficient (CC) has been observed to be a reliable indicator of success for structures with a resolution of 2.5 Å or better (Thorn & Sheldrick, 2013). Thus, for those MR trials with a correctly placed search model, a *SHELXE* correlation coefficient (CC) of at least 25 was used as an additional success criterion.

Due to the special characteristics of coiled-coils the generally accepted metrics of success do not always correctly indicate an actual solution (Thomas et al., 2015, 2020), so these cases required additional examination. At first, it was determined whether each coiled-coil case was solved or not by following the same procedure described in a previous *AMPLE* coiled-coil case study (Thomas et al., 2020). Accordingly, for each structure, the solution with the highest ranking *SHELXE* correlation coefficient was used for automated model building using *PHENIX AUTOBUILD* version 1.17 (Liebschner et al., 2019). This was performed using the *SHELXE* output build as the initial model, and it was tested both with and without the use of NCS for density modification. Successful solutions were then determined by *R*_free_ values below 0.45, the completeness of the model and a correlation coefficient between its *2F_o_ - F_c_* map and that of the deposited structure above 0.60. For the purpose of the data analysis required to compare the performance of the different ensembles in the new library, MR runs that met the success criteria used for the transmembrane and globular folds were only considered successful if the top solution was found to be a success using the above additional criteria.

All software used in the study corresponds with the CCP4 version 7.073: *PHASER* version 2.8.2, *REFMAC* version 5.8 and *SHELXE* version 2019/1. Calculation of correlation coefficients between electron density maps was performed using *phenix.get_cc_mtz_mtz* from the *PHENIX* suite version 1.17.

## 3. Results and Discussion

### 3.1 The new ensemble library solves more cases than the set of ideal helices

To assess the performance of single model ideal helices against their ensemble counterparts, a set of 111 structures were selected as described above. To test performance across different types of structures, this set was composed of transmembrane, globular and coiled-coils: 34, 31 and 46 structures respectively. Using *AMPLE’s* ideal helix and helical ensemble modes, a solution was attempted for each of these structures, using the original library of ideal helices and the members of the newly generated ensemble library as search models respectively.

For 46 out of the 111 structures of the dataset, at least one of the members of the original *AMPLE* ideal helix library was placed correctly by *PHASER* and *SHELXE* was able to successfully build a model out of this initial placement (Supplementary Table 5). The range of solutions achieved by these single model ideal helices is broad (Figure 3) with solutions up to resolutions of 2.6 Å (PDB code: 6I6B), 570 residues in the asymmetric unit (PDB code: 4FP4) and an eLLG as low as 26 (PDB code: 5ZLE). The number of solutions achieved when using the new library of ensembles rises to 61, an increase of 30 % on the total number of solutions achieved using ideal helices. Curiously, this increase was not uniform across the three fold types under study, with the highest increase being observed on transmembrane structures (40%), followed by globular (30%) and coiled-coils folds (25%). Additionally, the use of the new library of helical ensembles enabled *AMPLE* to solve increasingly challenging structures across the three folds, as is the case of 1D7M, a coiled-coil structure with a resolution of 2.7 Å and 404 residues in the asymmetric unit (ASU), the globular structure 5MQ8 which has 325 residues in the ASU and a resolution of 2.25 Å, or the case of the transmembrane structure 4RI2, with a resolution of 2.35 Å and 412 residues in the ASU (Figure 4). Encouragingly, this increase in the absolute number of solutions was achieved with no loss of prior solutions: all of the structures solved using single ideal helices were also solved with at least one member of the new ensemble library.

**Figure 3.**
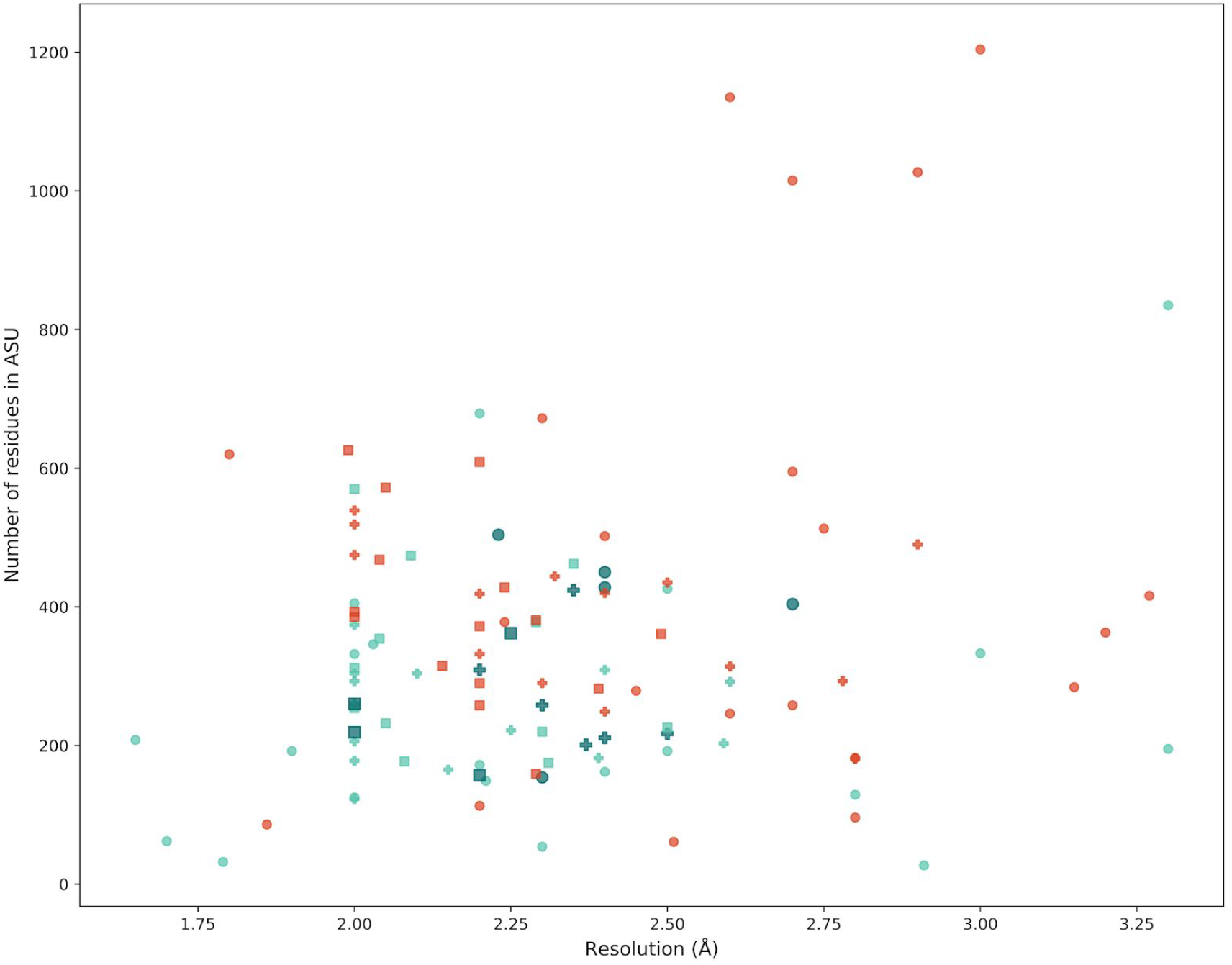
Distribution across different resolutions and number of residues in the asymmetric unit (ASU) of the structures solved using ideal helices and ensembles for the three different datasets. Each point represents a case of the dataset: transmembrane helical structures are represented with a cross, alpha globular structures with a square and coiled-coils with circles. Red points indicate a case that was solved by neither ideal helices nor ensembles. Light turquoise points represent a case solved by both ensembles and single models. Darker turquoise points indicate a case that could only be solved by ensembles. Full results can be seen at Supplementary Table 5.

**Figure 4.**
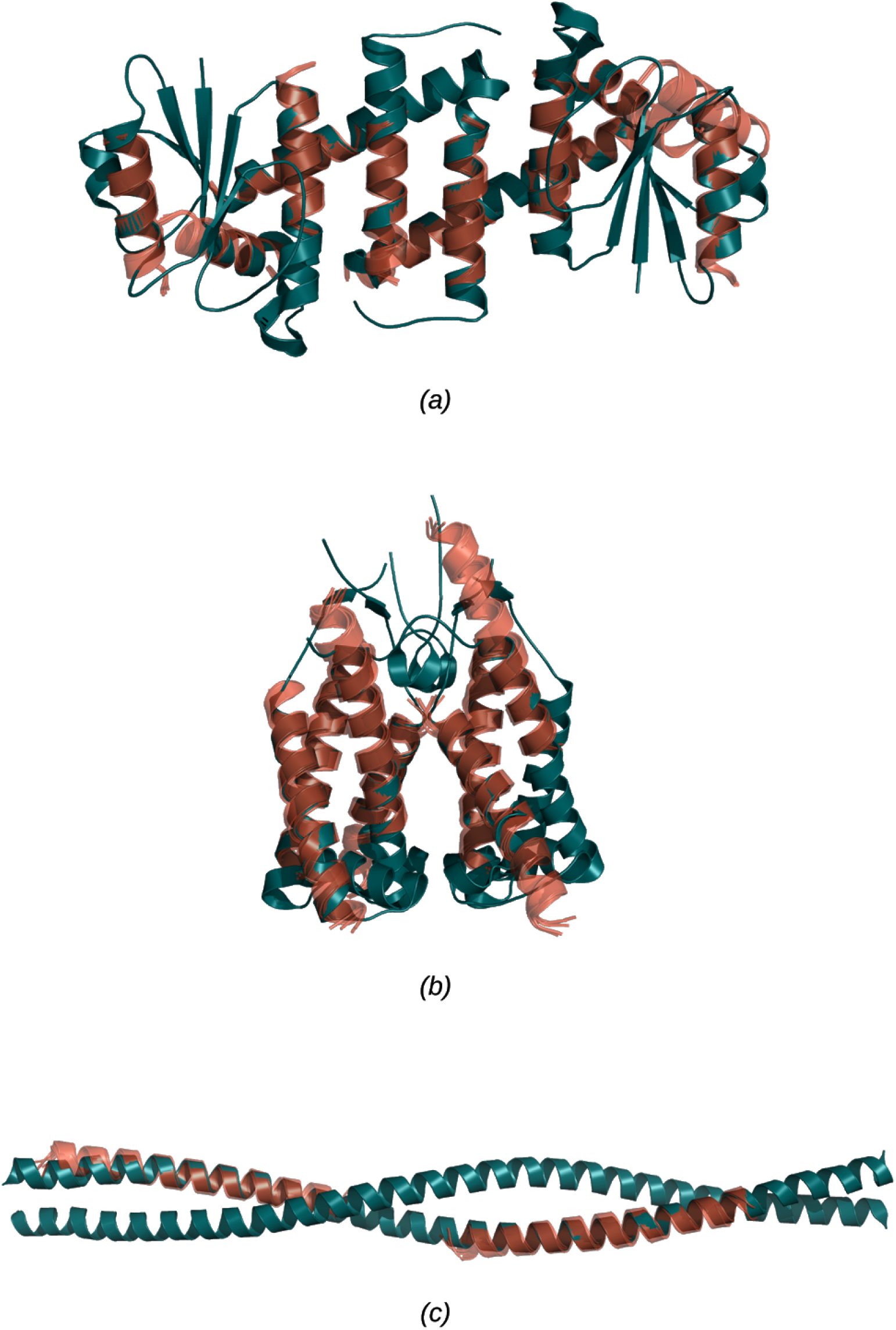
Examples of successful MR runs using members of the new library of helical ensembles. Figure A corresponds with the globular structure 5MQ8, with 12 placed copies of the 15 residue long helical ensemble. Figure B illustrates the solution of the transmembrane structure 4RI2, with 7 placed copies of the 25 residue long helical ensemble. Figure C shows the solution for the coiled-coil structure 1D7M, with 2 copies of the 35 residue long ensemble. In all three figures green chains correspond with the deposited crystal structure and the purple chains with the MR-placed ensembles (only the first model of the ensemble is shown).

*SHELXE* CC can be used as an excellent guide to correct MR solutions in most cases with a resolution better than 2.5 Å (Thomas et al., 2015, 2020; Thorn & Sheldrick, 2013). Encouragingly, 90% of those search models that were placed correctly according to our primary success criterion - a map CC of at least 0.2 - reported a *SHELXE* CC higher than 25. The high proportion of successful rebuilds achieved by *SHELXE* reflects the ability of its auto-tracing algorithm to correctly build the target’s main-chain with great accuracy. However, as is well known (Thomas et al., 2015, 2020; Thorn & Sheldrick, 2013) this metric loses accuracy in cases with poor resolution or for coiled-coils folds. In this regard, out of the 71 structures that scored *SHELXE* CC values above 25, 12 were identified as MR failures when the map CC was examined. The resolutions of these MR trials ranged between 1.67 Å and 3.2 Å, with an average of 2.2 Å. All corresponded to coiled-coil structures except for 4RYM, a transmembrane structure with a resolution of 2.8 Å.

In order to quantify the extent at which using the new ensembles extends the range of *AMPLE* solutions towards increasingly difficult cases, the eLLG for each of the solved structures was calculated using a 40 residue-long ideal helix as a search model: this eLLG can be used as an indicator of the overall difficulty of a case with lower values indicating greater difficulty. A comparison of the distribution of eLLG values between different categories of success was then made (Figure 5). Although the numbers are relatively small, differing patterns in the eLLG values of the additional solutions achieved by the ensembles in each of the three fold classes were observed. Both transmembrane and coiled-coil structures that could only be solved making use of the ensemble library have eLLGs within the bottom 25% of the base solutions obtained with ideal helices, an indication that using ensembles extends the range of solutions obtained using *AMPLE* ideal helix mode towards increasingly difficult cases. Curiously, this pattern was not observed in the globular dataset, where the structures that could only be solved by the members of the ensemble library present a high eLLG distribution, having most of them eLLG values within the top 25% of the structures solved with the original set of ideal helices. Therefore it appears that, in contrast with the other two fold types, rather than enabling *AMPLE* to solve increasingly difficult globular structures, using the new ensemble library yields solutions of those cases that could not be solved by ideal helices, despite having relatively high eLLGs.

**Figure 5.**
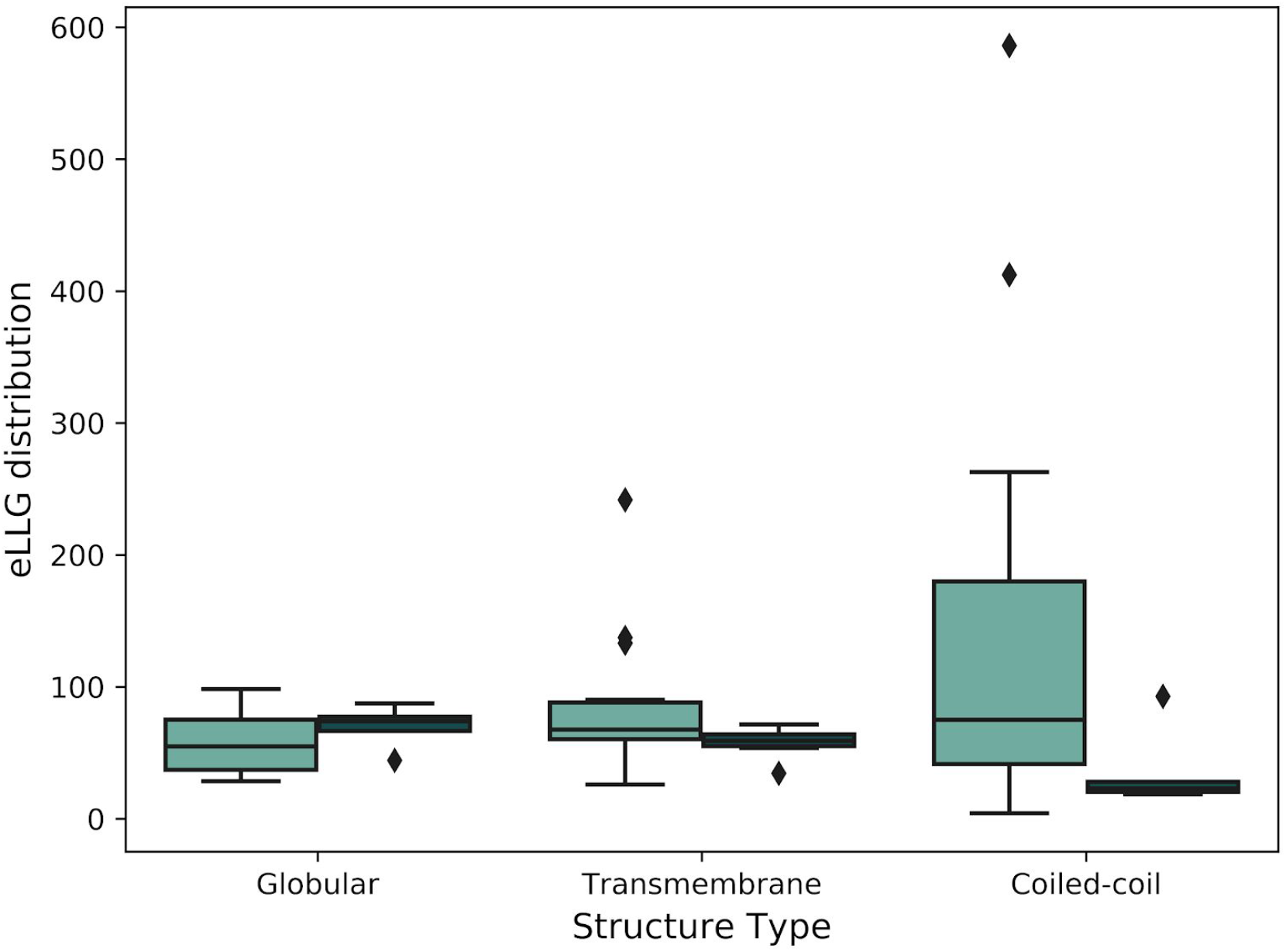
A box plot of the range of eLLGs for structures that were solved both using single model ideal helices and ensembles (light turquoise) and those that could only be solved by *AMPLE* when using members of the new ensemble library as search models (dark turquoise). Box limits indicate upper and higher quartiles, whiskers indicate upper and lower bounds and a horizontal line in the middle of the box plot represents the median. Outliers are depicted as a rhombi.

### 3.2 The optimal helical ensemble length varies with fold class

The original ideal helix library first implemented in *AMPLE* had eight ideal helices, with a size range starting at five residues and extending to 40 with a five-residue step (Thomas et al., 2017). In order to assess how the size of the search model affects the outcome of MR, search models were grouped into size-bins and the number of solved structures observed at each bin measured across the three datasets (Figure 6). Interestingly, the most successful search model size in the transmembrane dataset was 25 residues, 15 residues in the case of the globular structures and 30 residues for coiled-coils. These differences across the three datasets possibly reflect the different nature of these structures as coiled-coils tend to consist of very elongated helices, helical regions of globular proteins have a broad range of sizes and the thickness of most lipid bilayers can accommodate helices of approximately 20 to 30 residues (Hildebrand et al., 2004). It is also possible to observe that search models with only 10 residues or fewer were the least successful consistently across the three datasets, something especially noticeable in the case of transmembrane structures where no solutions were observed when using 5 residue-long helices.

**Figure 6.**
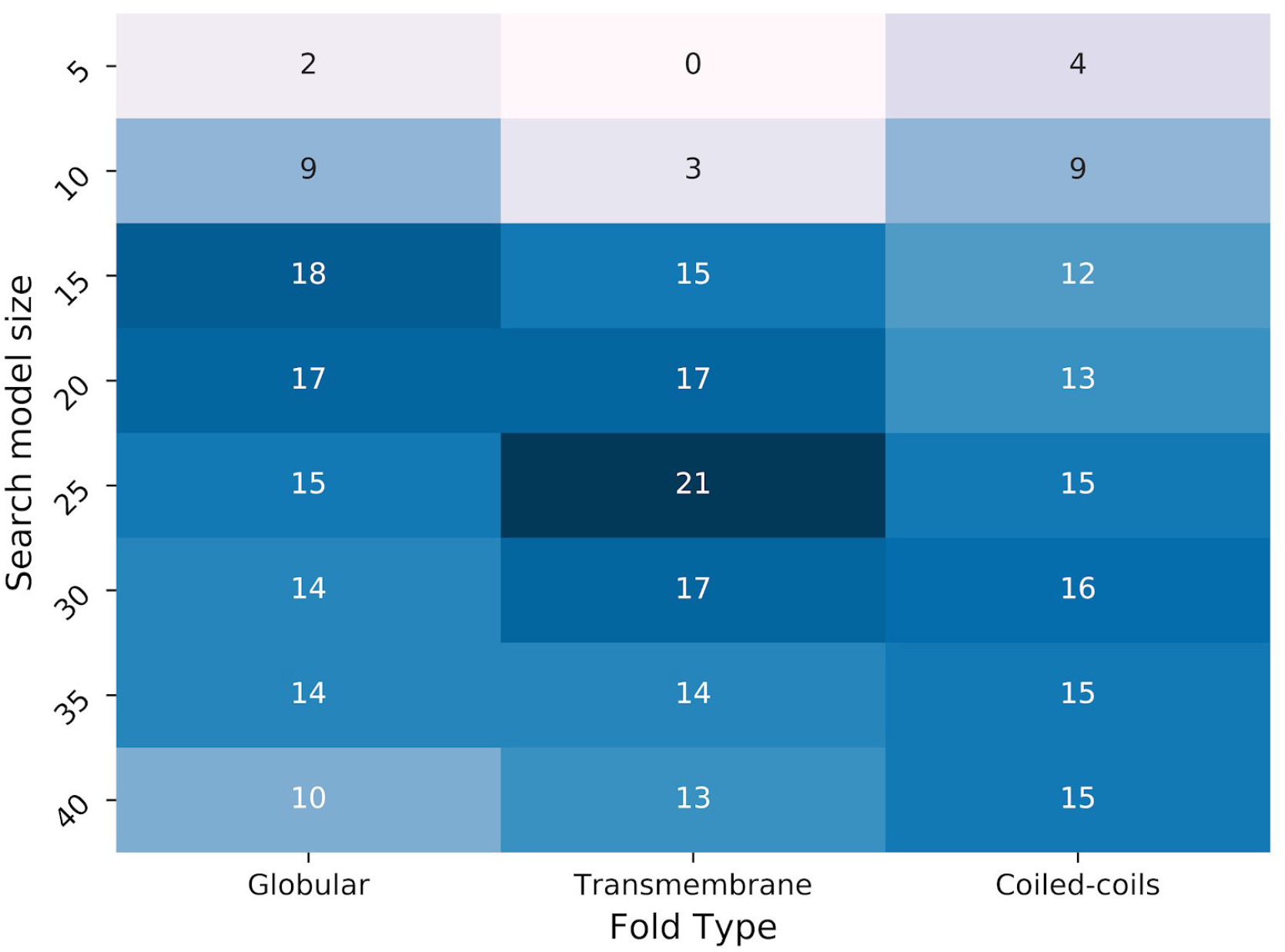
Analysis of the percentage of solved structures per ensemble size across the three fold types. Each cell represents the percentage of solutions that can be obtained using exclusively ensembles of each specific size, and have been colored accordingly in a blue scale from dark blue -higher percentage- to light blue -lower percentage-.

### 3.3 Ensemble heterogeneity does not affect search model effectiveness

For each possible combination of helix size and B-factor treatment, both a low divergence homogeneous ensemble and a more heterogeneous ensemble with high variability between its models were created. In order to assess if the structural similarity between the different models that comprise the ensemble has an effect on the effectiveness of the search model, results with these two versions of ensembles were analysed (Figure 7). This revealed no significant differences in the number of solutions obtained using homogeneous ensembles and their heterogeneous counterparts in any of the three structural folds, an indication that search model effectiveness is not affected by changes in the level of ensemble heterogeneity within the range of values being tested in this study.

**Figure 7.**
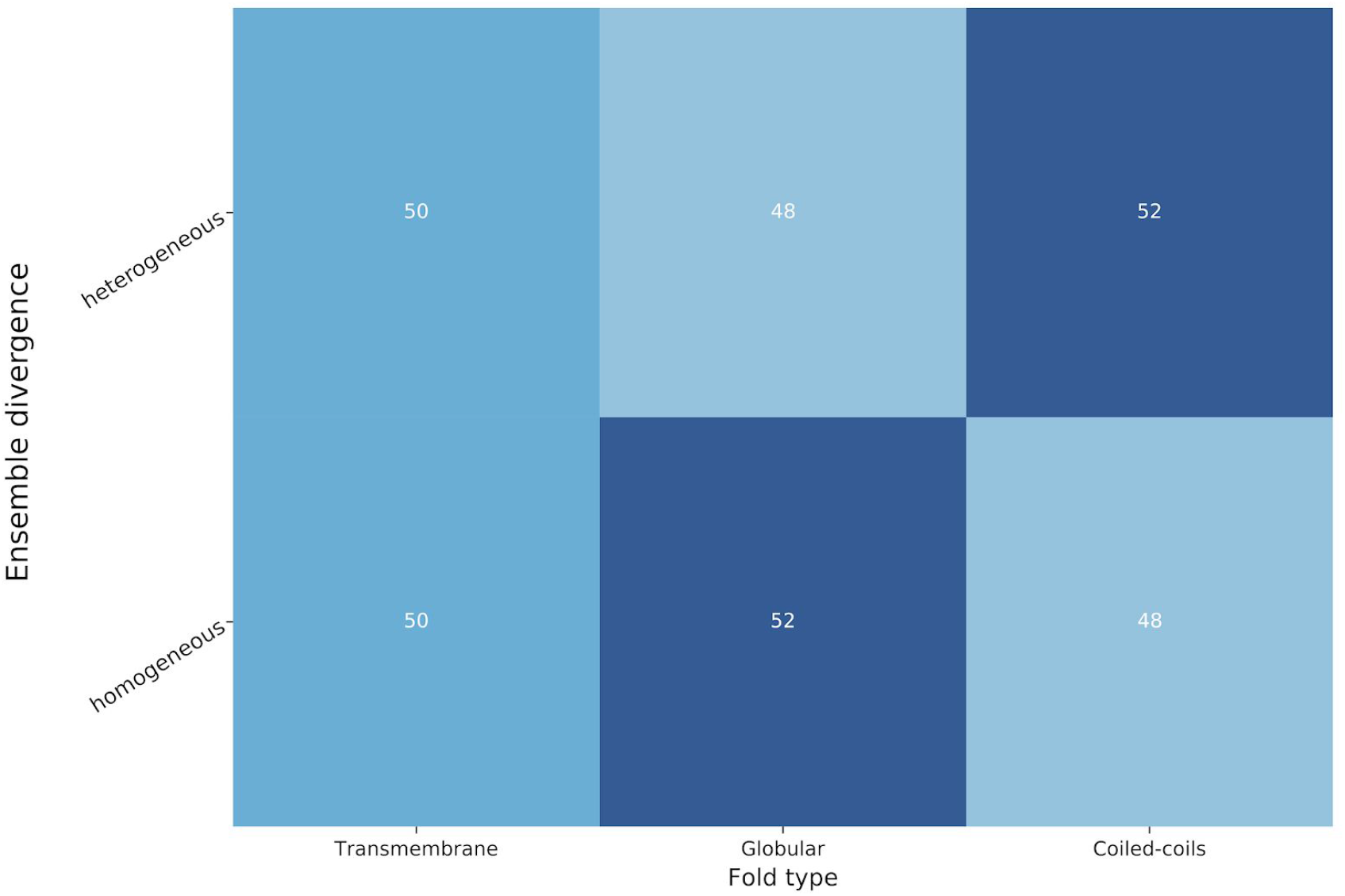
Analysis of the percentage of solved structures per ensemble divergence across the three fold types. Each cell represents the percentage of solutions that can be obtained using exclusively ensembles of each specific heterogeneity, and have been colored accordingly in a blue scale from dark blue -higher percentage- to light blue -lower percentage-.

Despite not having observed significant effects of the ensemble heterogeneity in the total number of solved structures, a comparison of the effectiveness of homogeneous and heterogeneous search models revealed that the success ratio of these two types of ensembles varies across different resolution ranges (Supplementary Figure 1). Interestingly, homogeneous ensembles appeared to be more successful for structures with resolutions both worse than 2.75 Å and better than 2.00 Å, while it was not possible to appreciate any tendency for the rest of structures within the 2.00-2.75 Å range.

**Supplementary Figure 1.**
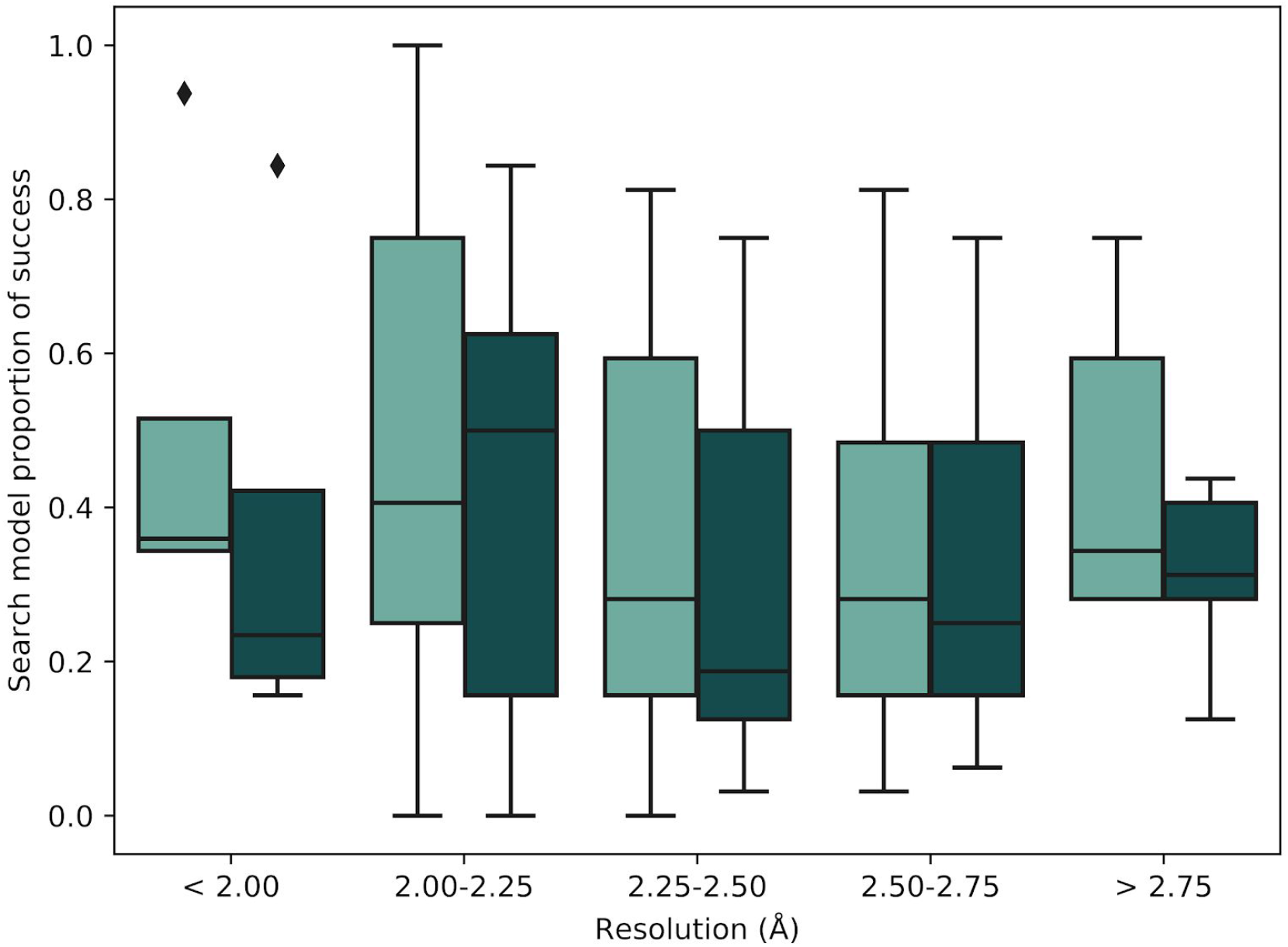
Distribution of the proportion of search model successes across different resolution ranges between homogeneous ensembles (light turquoise boxes) and heterogeneous ensembles (dark turquoise boxes). Box limits indicate upper and higher quartiles, whiskers indicate upper and lower bounds and a horizontal line in the middle of the box plot represents the median. Outliers are depicted as a rhombi.

### 3.4 A minimal set of ensembles can be used without any solution loss

In order to assess if there was any level of redundancy among the different ensemble B-factor treatments in the new library, the structures solved by each ensemble were gathered and the common solutions across different ensemble preparations compared (Figure 8). It is interesting to observe that the four B-factor treatments shared 52 out of the 61 solutions achieved by the ensemble library, indicating a high level of similarity in the performance of these ensemble treatments. Interestingly, this similarity was also observed across different resolution ranges, as no evident differences in performance were revealed after performing an analysis of the search model effectiveness across different resolutions (Supplementary Figure 2). Nevertheless, not all ensembles contributed in the same way to solve the additional targets that could not be solved using the original *AMPLE* library of single model ideal helices. Thus, only 5 out of the 17 newly solved targets succeeded with ensembles from all four four B-factor treatments, indicating the importance of screening the broad spectrum of ensemble treatments to gain elusive solutions that could not be solved with the original ideal helices. However, it is important to note that a reduced set of ensembles could still be created if required without losing any of those additional solutions, as search models with B-factor treatments 2 and 3 cover the full range of extra solutions obtained using ensembles.

**Figure 8.**
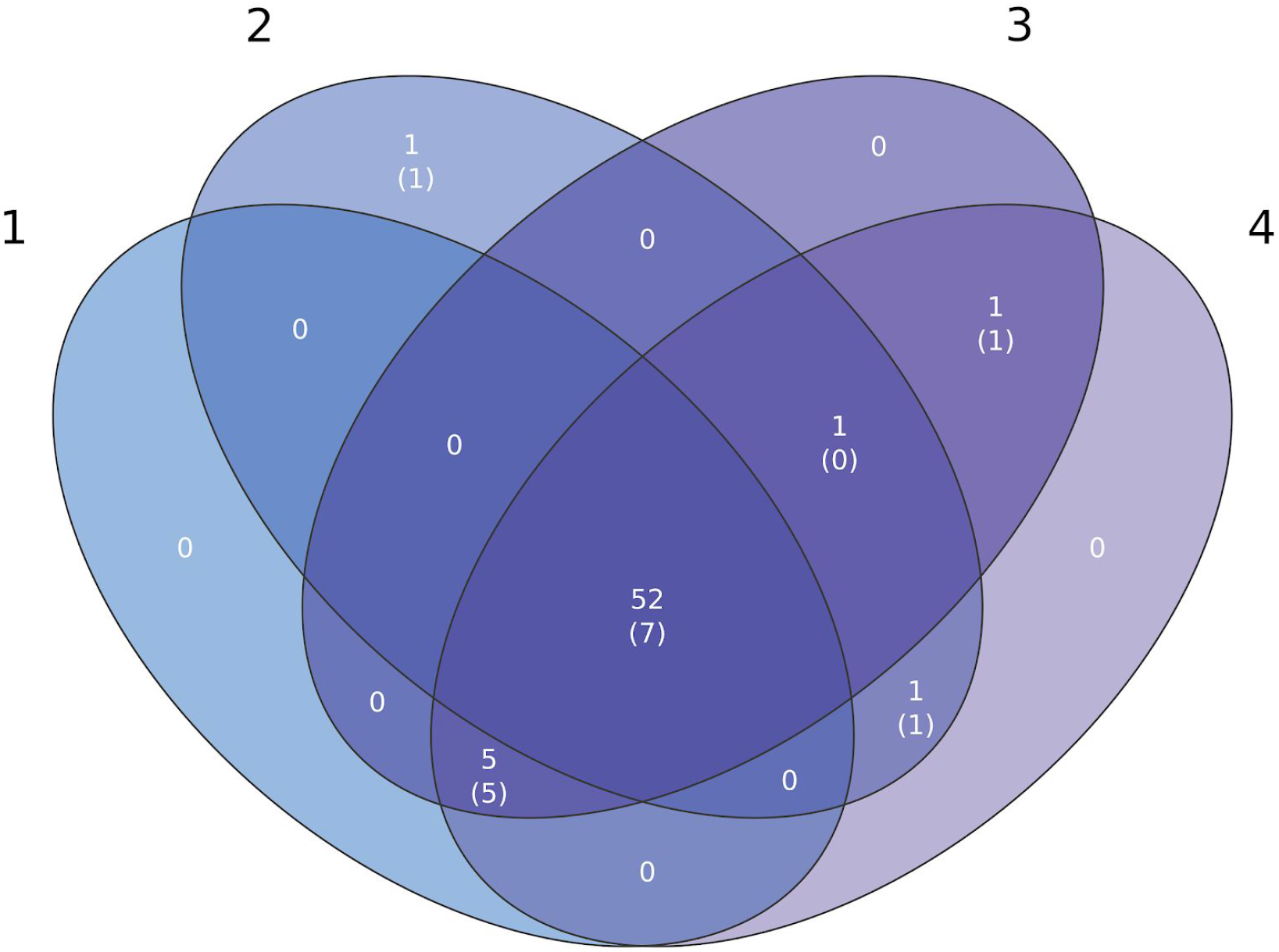
Venn diagram with the number of solved structures achieved with each B-factor treatment. Black numbers indicate B-factor treatment (see text for details), the total number of solutions is indicated with white inner numbers, from which solutions that could only be achieved with members of the ensemble library and not the original *AMPLE* ideal helices are between brackets.

**Supplementary Figure 2.**
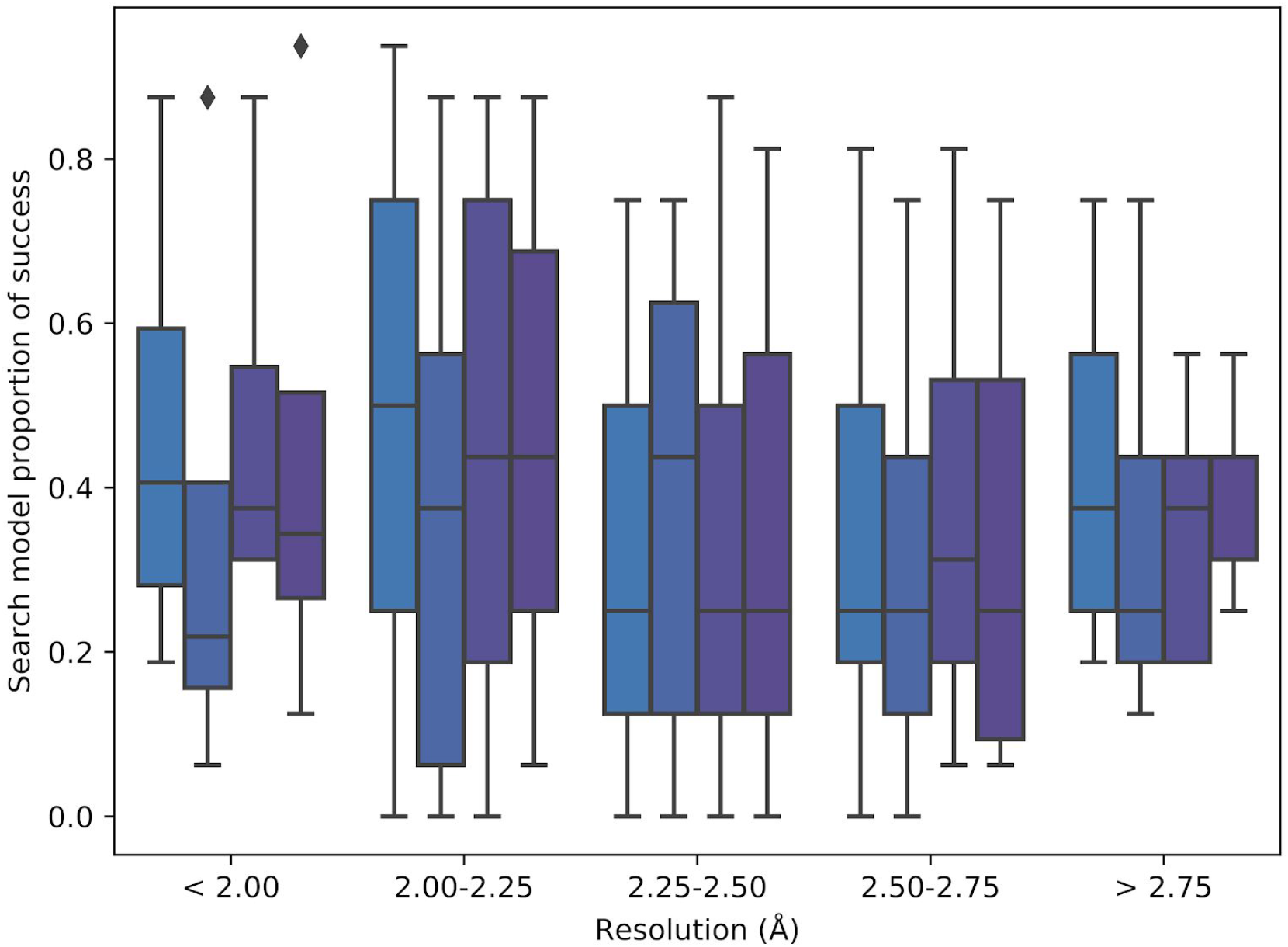
Distribution of the proportion of search model successes across different resolution ranges between different bfactor treatments -from blue to purple-: treatment 1, 2, 3 and 4. Box limits indicate upper and higher quartiles, whiskers indicate upper and lower bounds and a horizontal line in the middle of the box plot represents the median. Outliers are depicted as a rhombi.

### 3.5 A minimal set of ensembles yields solutions faster than the original library of ideal helices

The new library of ensembles consists of 64 members, which represents an eight-fold increase in the number of search models over the original library of ideal helices. In order to assess whether this increase in the library size translates into an increase in the computing time consumed by *AMPLE*, we compared the timings of both approaches in those cases where at least one of the members of both libraries yielded a solution. To do this, we registered the time required to reach a solution by using previously recorded timings of MR runs and simulating four different MR strategies. The first simulation (simulation 1) corresponded with the time spent by AMPLE using the new library of helical ensembles, while the second simulation with the use of the original set of ideal helices (simulation 2). In these first two simulations, search models were ordered by increasing chain size, and no specific priority was set among the different B-factor treatments in the new library. Additionally, we were also interested in extending this comparison to a minimal subset of ensembles, which was created using the observations made about the performance of the different ensembles in the previous section. Since ensemble heterogeneity had no major effects on search model success, only the homogeneous ensembles were taken into the minimal subset (Figure 7). Additionally, since 5 and 10 residue long ensembles were observed to be the least successful (Figure 6), helical ensembles of this size were removed from the third simulation. Regarding B-factor treatments, results showed that there is no solution loss when only using ensembles with B-factor treatments 2 and 3 (Figure 8), which were taken into the minimal subset of ensembles. All these changes resulted in the creation of a subset of 12 ensembles, which were trialled on a third and fourth simulations. These two last simulations differed on the order at which these ensembles were trialled. On the third simulation, the search model sorting was determined by the overall success rates previously observed for each ensemble size, independently of the fold class, and the search models with the most successful sizes were trialled first (simulation 3). Finally, since we observed that the optimal ensemble size varies across fold types (Figure 6), we implemented a fold-specific ordering on a fourth simulation (simulation 4).

Interestingly, the increase in the number of search models in the new ensemble library translates into an increase in the time necessary before a solution is found in simulation 2 when compared with the time required by *AMPLE* when the original ideal helices are used in simulation 1 (Figure 9). Nevertheless, when using the minimal subset of ensembles, simulations 3 and 4 show that *AMPLE* was able not only to match the time performance of the single model library, but to yield a solution faster across most targets in the dataset. This reveals a further advantage of using the new ensemble library: it not only achieves more solutions than the original ideal helix library but also reaches a solution faster in those cases where the solution can be found by the original library. Interestingly, despite having observed that ensemble sizes have different success rates depending on the fold class of the unknown structure, we observed only minor improvements in the time elapsed before a solution is found when a fold-specific search model ordering is used, as differences between simulations 3 and 4 are minimal in most cases.

**Figure 9.**
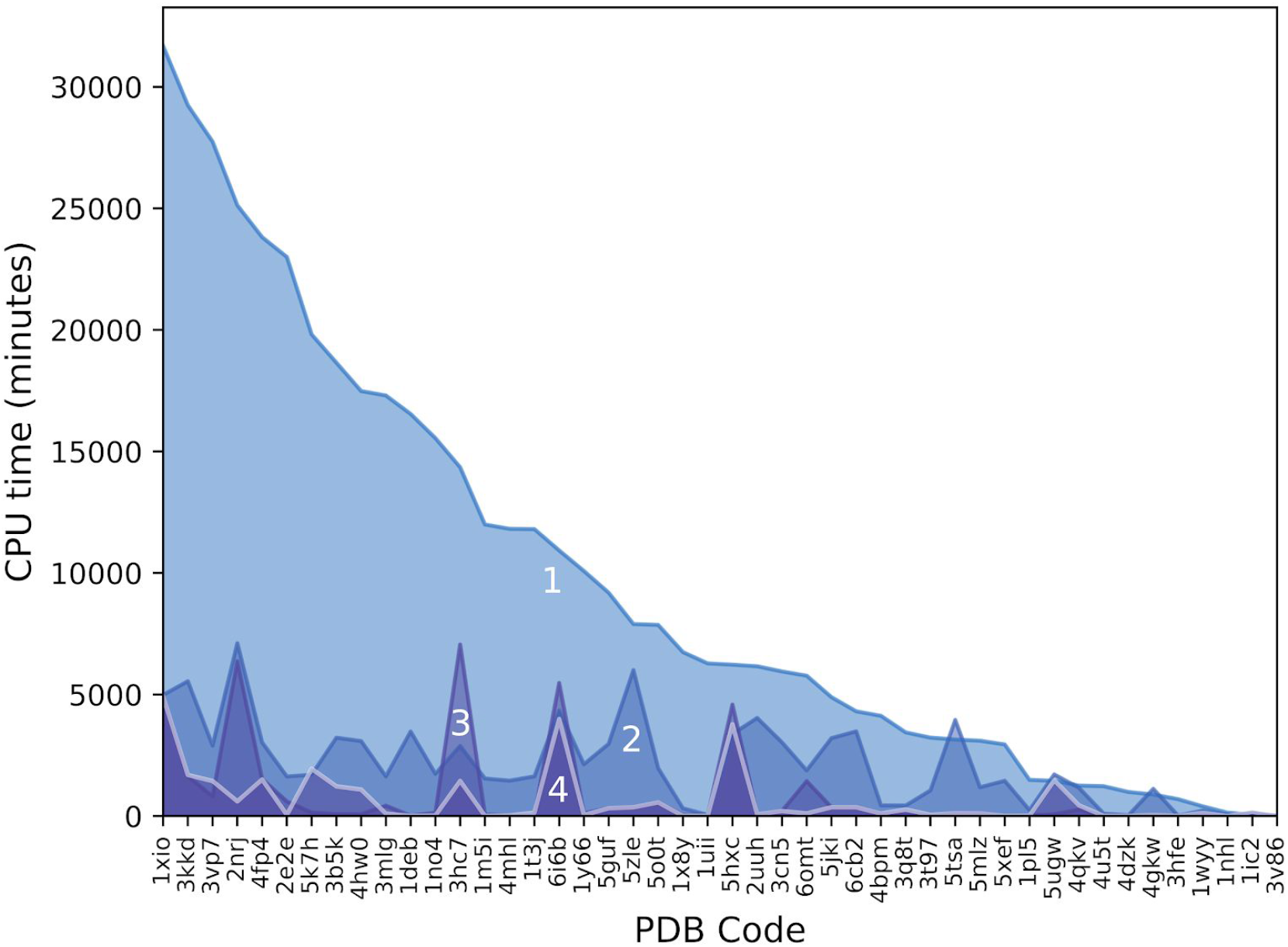
Elapsed time -obtained by simulations based on job timings- before *AMPLE* reaches the first solution using as search models the members of the new ensemble library (1), the original library of ideal helices (2), and a minimal subset of the new ensemble library using fold-independent search model sorting (3) and fold-specific ordering (4).

## 4. Conclusion

Here we have presented a new take on the concept of using helices as search models by making use of a new set of 64 helical ensembles. The use of the new library of ensembles resulted in a 30% increase in the number of solutions compared to a library of ideal helices, which was consistent across the three folds under study. Having observed no solution loss when using ensembles, we strongly encourage the use of this new library of search models as an alternative to ideal helices. These findings agree with observations made by previous studies, where clustering several search models into ensembles outperformed the individual use of these models (Keegan et al., 2018; Rigden et al., 2002; Simpkin et al., 2020) as their structural variability can be used to statistically weight sets of structure factors (Read, 2001). We also observed that some solutions required the use of ensembles modified with a series of B-factor treatments reflecting the structural variability across the models of the ensemble. These findings stress the importance of B-factors on *PHASER’s* algorithm, as the structure factors are computed based on the normalised B-factors of the atoms in the supplied search model, before actual MR operations begin.

Based on observations made about the efficacy of our new search models, we have been able to create a minimal subset of 12 ensembles without any solution loss compared with the full size library, revealing that our new approach does not require intensive use of computational resources to improve upon results obtained with previous methods. Thus, we believe that usage of our minimal subset of helical ensembles can be an alternative MR route to more time-consuming approaches required when no homologous structures can be found. In contrast with some of these approaches that rely on precise tertiary structure predictions to produce suitable search models (Keegan et al., 2015; Rigden et al., 2018; Simkovic et al., 2016), our take on fragment-based MR only requires the presence of alpha helices within the unknown structure, which is known to be the most reliably predicted regular secondary structure element (Cuff & Barton, 2000).

We expect that our new set of ensembles will also be able to provide assistance on other commonly observed limitations of fragment-based MR, especially in regards to the completion of partial MR solutions, which are commonly observed on this approach (Millán et al., 2020). Since the search models employed in fragment MR normally constitute a small portion of all the residues present in the unsolved structure, model building and chain tracing can be unsuccessful despite having correctly placed the search model. Our unpublished preliminary work has revealed that after successfully placing a small fragment with high local structural similarity to the unknown structure, it is possible to successfully complete the partial MR solution using our new helical ensembles as search models, where ideal helices deployed for the same purpose failed. Then, the placement of these additional search models together with the initial fragment provides enough signal to solve the structure through model building and density modification.

